# A first draft genome of Holm oak (*Quercus ilex* L.), the most representative species of the Mediterranean forest and the Spanish agrosilvopastoral ecosystem “*dehesa”*

**DOI:** 10.1101/2022.10.09.511480

**Authors:** María-Dolores Rey, Mónica Labella-Ortega, Víctor M. Guerrero-Sánchez, Rômulo Carleial, María Ángeles Castillejo, Antonio Rodríguez-Franco, Richard G. Buggs, Valentino Ruggieri, Jesús V. Jorrín-Novo

## Abstract

The holm oak (*Quercus ilex* L.) is the most representative species of the Mediterranean Basin and the agrosilvopastoral Spanish “*dehesa*” ecosystem. Being part of our life, culture, and subsistence since ancient times, it has great environmental and economic importance. More recently, there has been a renewed interest in using the *Q. ilex* acorn as a functional food due to its nutritional and nutraceutical properties. However, the holm oak and its related ecosystems are threatened by different factors, with oak decline syndrome and climate change being the most worrying on the short and medium term. Breeding programs informed by selection of elite genotypes seems to be the only plausible biotechnological solution to rescue populations under threat. To achieve this and other downstream analyses, we need a high-quality *Q. ilex* reference genome. Here, we introduce the first draft genome assembly of *Q. ilex* using long-read sequencing (PacBio). The assembled nuclear haploid genome has 530 contigs totaling 842.2 Mbp (N50 = 3.3 Mbp), of which 448.7 Mb (53%) are repetitive sequences. We annotated 39,443 protein-coding genes and Benchmarking Universal Single-Copy Orthologs analysis detected 412 out of 425 expected complete and single-copy genes (94.80%) within the *Q. ilex* genome. The chloroplast genome size was 142.3 Kbp with 149 protein-coding genes successfully annotated. This first draft should allow for the validation of - omics data as well as the identification and functional annotation of genes related to phenotypes of interest such as those associated to resilience against oak decline syndrome and climate change, higher acorn productivity and nutraceutical value.

## Background

Holm oak (*Quercus ilex* L.) is included in the 640 recorded species in the “Plants of the World Online” database within the genus *Quercus* (de Rigo and Caudullo, 2016; Kew Royal Botanic Gardens; https://powo.science.kew.org/taxon/urn:lsid:ipni.org:names:325819-2#bibliography). It is one of the most important clades of woody angiosperms in the Northern hemisphere in terms of species diversity, ecological dominance, and economic value (Backs and Ashley, 2021), and has been part of our life, culture, and subsistence since ancient times (Leroy et al., 2020). Moreover, it is a priority species for afforestation due to its high adaptability to semi-arid climate conditions, drought tolerance (Barbeta and Peñuelas, 2016), and plastic response to varying edaphic conditions (Laureano et al., 2016).

The holm oak gives personality to the *dehesa*, an agrosilvopastoral ecosystem of great environmental and economic importance, especially in rural areas (Plieninger et al., 2021). The most remarkable economic benefit is related to its fruit, the acorn. For example, costumers tended to prefer the flavour of ham from acorn-fed black Iberian pigs (*Sus scrofa domesticus*) (Diaz-Caro et al., 2019). Recently, there has been a renewed interest for *Quercus* spp., to be used a source of functional food due to the increasing demand for alternative plant species, including forest trees, for dietary diversification, sustainable food production, and nutrition (Vinha et al., 2016; Lopez Hidalgo et al., 2021).

Currently, *Q. ilex* is seriously threatened by several anthropogenic and environmental factors including high average population age, overexploitation coupled with poor regeneration capacity, inappropriate livestock management, and biotic (e.g., *Phytophthora cinnamomi*) and abiotic stresses (e.g., extreme temperatures and extended drought periods). Together, these factors likely contribute to the so-called holm oak decline syndrome, which has been causing an increase in tree mortality and has seriously damaged the *dehesa* (Brasier, 1992; Camilo-Alves et al., 2013; Ruiz-Gómez et al., 2018; San-Eufrasio et al., 2021).

The main priority for *Q. ilex* conservation and exploitation is the establishment of breeding programs aimed at increasing production and resilience to the decline syndrome and climate change (Cortés et al., 2020). These programs will contribute to the proper and sustainable management, conservation, and exploitation of *Q. ilex* and related ecosystems (Plomión et al., 2016). A better understanding of this species at the biochemistry, molecular biology, and biotechnology levels will facilitate these breeding programs, as it has occurred with many herbaceous and other woody species (Muthuramalingam et al., 2022). As with many long-lived, non-domesticated and allogamous species, the most plausible breeding strategy for *Q. ilex* is the use of genome-wide association studies (GWAS) to identify elite genotypes associated with phenotypes of interest (e.g., high acorn production with desirable quality traits related to nutritional values, or resilience to adverse biotic and abiotic stresses; Maldonado-Alconada et al., 2022). So far, the number of publications focusing on the molecular aspects of *Q. ilex* biology has been quite limited due to its challenging biological characteristics and its recalcitrance as an experimental system (Rey et al., 2019; Escandón et al., 2021; Maldonado-Alconada et al., 2022). However, despite these difficulties, our group has to some extent characterized the genetic structure and diversity of *Q. ilex* using DNA markers, microsatellite, transcriptomics, proteomics, and metabolomics (Valero-Galván et al., 2011; Fernández i Marti et 2018; Lopez-Hidalgo et al., 2022; Maldonado-Alconada et al., 2022). These different approaches have allowed us to study seed germination responses to biotic and abiotic stresses and their interaction, and also the integration between molecular and morphometric data (Escandón et al. 2021). However, our research has so far been constrained by the lack of a *Q. ilex* reference genome, so the picture provided by these studies is still incomplete. The sequencing of the *Q. ilex* genome will add to the list of many recently published *Quercus* reference genomes (Plomión et al., 2016, 2018; Sork et al., 2016, 2022; Ramos et al., 2018; Han et al., 2022; Zhou et al., 2022, Ai et al., 2022), and will allow us to interpret *Q. ilex* - omics data in terms of its own gene products rather than relying on orthologs from other *Quercus* species. This should open new research avenues related to the genetic make-up of this species, the identification of novel genes, allelic variants, SNPs, epigenetic marks, and will allow for studies on broader topics such as phylogenetics, hybridization and introgression patterns, and breeding programs informed by GWAS and genome prediction (Plomión et al., 2018; Stocks et al., 2019; Sork et al., 2022).

Many recently reported *de novo* genome assemblies have been carried out using next-generation short-read sequencing technologies such as Illumina or 454 platforms (Zhao et al., 2017; Xia et al., 2017), which tend to produce very fragmented genomes. Pacific BioSciences (PacBio) has developed a third-generation sequencing technology to assemble large and complex plant genomes producing tens of thousands long individual reads (up to ~40 kb) (Roberts et al., 2013; Paajanen et al., 2019). Here, we generate a high-quality draft genome assembly for *Q. ilex* (NCBI’s BioProject repository, ID: PRJNA687489) using the single-molecule real-time (SMRT) sequencing technology from PacBio. We reveal genomic features of *Q. ilex*, including nuclear and chloroplast sequences, repeat sequences, and gene annotations.

## Data description

### Genomic DNA extraction, sequencing, and genome size estimation

Leaves were collected from a mature, healthy *Q. ilex* tree (Figure 1) located in Aldea de Cuenca, Fuente Obejuna, Córdoba, Andalusia, Spain (38° 19’ 46” N, 5° 33’ 15” W) in November 2019. Upon arrival at the laboratory, leaves were washed with 2% sodium hypochlorite and abundantly rinsed with distilled water. For long-read sequencing, extraction of high molecular weight genomic DNA from leaves was carried out using the modified cetyl trimethyl ammonium bromide (CTAB) method (Murray and Thomson, 1980) with slight modifications. Briefly, 4 gr of fresh leaves were macerated in liquid nitrogen. Since *Q. ilex* is a species rich in phenolic compounds, we added 1 gr of polyvinylpolypyrrolidone (PVPP) per 20 mL of CTAB. Once extracted, DNA was treated with RNAse (10 mg/mL, Thermo Fisher Scientific Inc., Waltham, MA, USA) for 1h 30 mins at 37ºC, and the quality and concentration were determined using a 1% agarose gel electrophoresis and a Qubit version 3.0 Fluorometer (Thermo Fisher Scientific Inc., Waltham, MA, USA), respectively.

**Figure 1:**
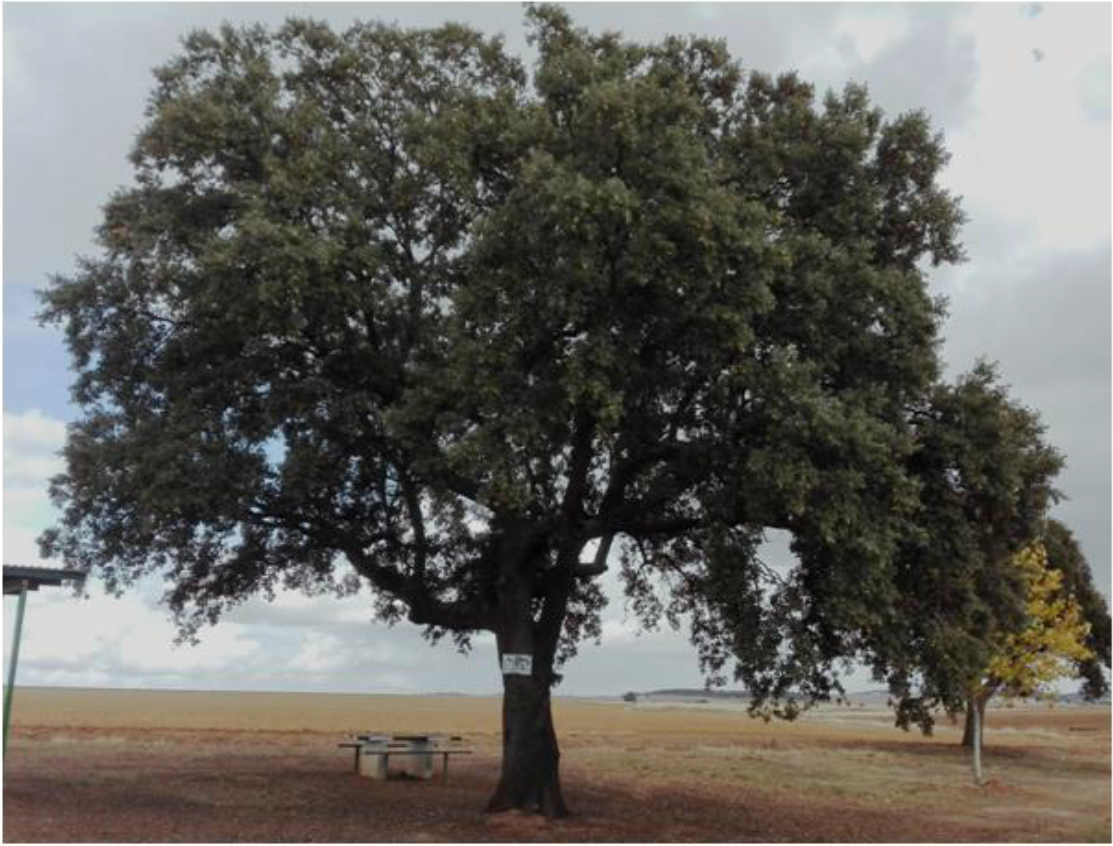
The sequenced individual used to produce the first *Q. ilex* draft genome. The tree is located in Aldea de Cuenca, Fuente Obejuna, Cordoba, Andalusia, Spain (UTM 30S 276751 4245466 datum ETRS89).

In the genus *Quercus*, both *Q. ilex* and *Q. suber* are sympatric in the Mediterranean area and hybrids have been identified in natural populations (Burgarella et al., 2009; López de Heredia et al., 2018). Previous studies based on interspecific genetic markers (alloenzymes, chloroplasts and nuclear DNA) have reported nuclear and cytoplasmic genome exchanges between both *Quercus* species (Toumi and Lumaret, 1998; Soto et al., 2007; López de Heredia et al., 2020). To determine whether the target individual (Figure 1) was a pure *Q. ilex*, we compared its genotype with genotypes from other nine *Q. ilex* and ten *Q. suber* mature individuals using 20 microsatellite markers as previously described in Fernández i Marti et al., (2018). PCR amplifications were conducted in a T100™ Thermal cycler (Biorad, Hercules, CA, USA) and PCR products were analyzed on an ABI Prism 310 capillary electrophoresis system (Applied Biosystems, Foster City, CA, USA). Size alignment and quality control were analyzed using GeneMapper version 3.7 (Applied Biosystems, Foster City, CA, USA). We performed a Principal Component Analysis (PCA) including all 20 microsatellite markers using the R package Adegenet (Jombart, 2008). *Q. ilex* and *Q. suber* individuals clustered at different sides of the PC1 axis (Supplementary Figure 1), indicating the purity of the target individual.

**Supplementary Figure 1.**
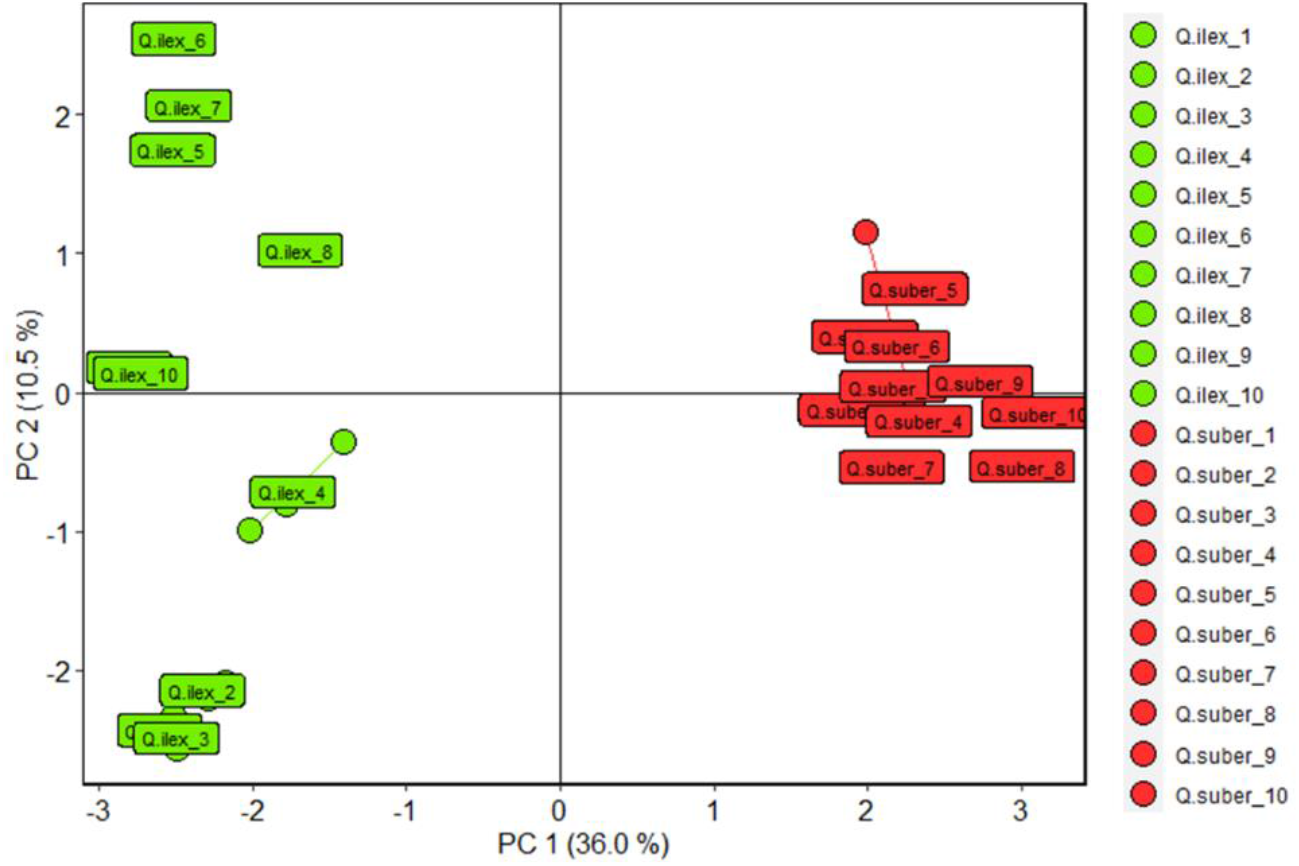
Principal component analysis of *Q. ilex* and *Q. suber* samples using 20 chloroplast and nuclear SSR markers. The analysis of the SSRs was conducted on ten biological replicates per each *Quercus* species (n = 20). The *Q. ilex* individual used for genome assembly is shown as *Q. ilex* 3.

The target DNA sample was used to construct a PacBio SMRT sequencing library at DNA Sequencing & Genotyping Center at the University of Delaware. Purified High Molecular Weight (HMW) DNA was fragmented to approximately 20 kb using Megaruptor version 3.0 (Diagenode, Denville, NJ, USA). 10 μg of purified HMW DNA was used as template to construct a SMRTbell HiFi library using Express Template Prep version 2.0 (Pacific Biosciences, Menlo Park, CA, USA) as per the manufacturer’s protocol. The quality of the HiFi library was assessed using the Qubit 3.0 Fluorometer (Thermo Fisher Scientific Inc., Waltham, MA, USA) and the Agilent Femto Pulse System (Agilent Technologies, Santa Clara, CA, USA). The HiFi library was run on two Sequel IIe system 8M SMRT cells using sequencing chemistry version 3.0 with 4 h pre-extension and 30 h movie time. Two runs of HiFi PacBio generated reads covering 64.3X of the genome assuming an a priori genome size of 850 Mbp. Although reads generated by HiFi PacBio are usually high quality, we performed an additional rough evaluation of nucleotide (A, C, G, and T) distribution to eliminate any remaining low-quality reads. Reads showing a skewed nucleotide distribution (approximately 1%) were removed.

### *De novo Q. ilex* genome assembly

Filtered reads were used as input for a *de novo* assembly approach undertaken using NextdeNovo version 2.5.0 software (key parameters: seed_depth=45, nextgraph: -a1) (https://github.com/Nextomics/NextDenovo). The software was run in “hifi” assembly mode, which skips the read correction step and produces a consensus of corrected reads using a string graph algorithm. The whole workflow is showed in Supplementary Figure 2.

**Supplementary Figure 2.**
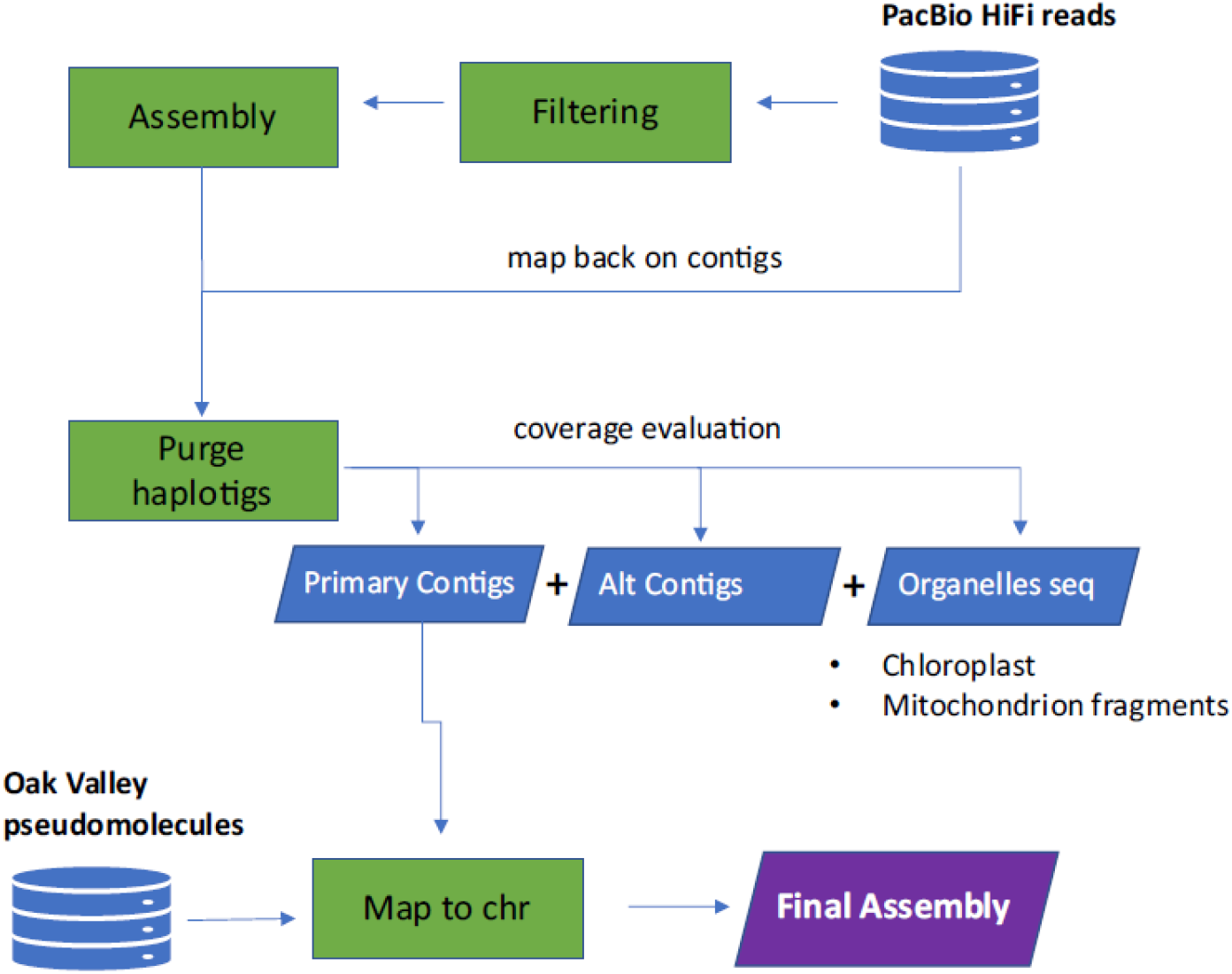
The *Q. ilex* genome assembly workflow.

We assembled 1,166 contigs totaling a genome size of 1,000,594,563 bp (Table 1). The N50 and the average contigs size were 2.63 and 0.85 Mbp, respectively. Genome size was higher than expected since assembly at this step includes primary, secondary, and organelle (chloroplast and mitochondria) contigs.

**Table 1.**
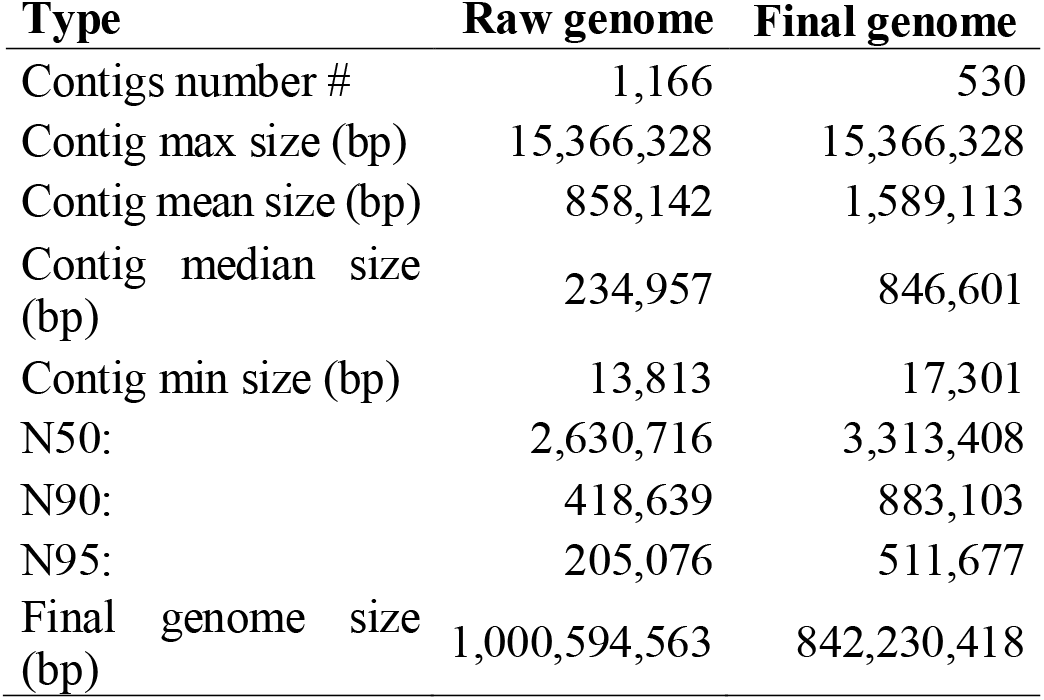
Raw and final genome assembly statistics

| Type | Raw genome | Final genome |
| --- | --- | --- |
| Contigs number # | 1,166 | 530 |
| Contig max size (bp) | 15,366,328 | 15,366,328 |
| Contig mean size (bp) | 858,142 | 1,589,113 |
| Contig median size (bp) | 234,957 | 846,601 |
| Contig min size (bp) | 13,813 | 17,301 |
| N50: | 2,630,716 | 3,313,408 |
| N90: | 418,639 | 883,103 |
| N95: | 205,076 | 511,677 |
| Final genome size (bp) | 1,000,594,563 | 842,230,418 |

Regions of very high heterozygosity can lead to problems during genome assembly (Kajitani et al., 2014). For example, once a pair of allelic sequences exceeds a certain threshold of nucleotide diversity, assembler may assemble these regions as separate contigs, rather than the expected single haplotype-fused contig, resulting in a significantly larger genome size. The *Quercus* genus is known to contain species with high heterozygosity (Sork et al., 2016; Ramos et al., 2018; Han et al., 2022, Ai et al. 2022), and the same was true for *Q. ilex* (1.93%) (Table 2).

**Table 2:** Comparison of summary statistics for genome assembly among *Quercus* species.

| Species | Genome size (Mpb/1C) | Heterozygosity (%) | Repeat elements (% of genome) | Protein coding genes (n) | Study |
| --- | --- | --- | --- | --- | --- |
| *Q. ilex* | 842.2 | 1.93 | 53.3 | 39,443 | This study |
| *Q. robur* | 814.3 | - | 53.3 | 25,808 | Plomión et al., 2018 |
| *Q. lobata* | 845.9 | 1.25 | 54.4 | 38,373 | Sork et al., 2016 |
| *Q. suber* | 953.5 | 1.62 | 12.0 | 33,658 | Ramos et al., 2018 |
| *Q. variabilis* | 787.0 | 2.15 | 67.6 | 32,466 | Han et al., 2022 |
| *Q. gilva* | 889.8 | 1.16 | 57.6 | 36,442 | Zhou et al., 2022 |
| *Q. mongolica* | 809.8 | 1.09 | 53.8 | 36,553 | Ai et al., 2022 |

To solve assembly issues, contigs were separated into primary contigs and associated secondaries contigs using the Purge_haplotigs pipeline version 1.1.1 (key parameter: default) (https://bitbucket.org/mroachawri/purge:haplotigs/src/master) (Roach et al., 2018). When plotting read coverage on the raw genome assembly contigs, we found a ‘haploid’ and a ‘diploid’ peak (Supplementary Figure 3). In regions with homologous contigs (e.g heterozygous regions) in the haploid assembly, reads were split between the two contigs resulting in approximately half the read-depth for those contigs. Based on the plot distribution, specific cut-offs were set to separate and flag the low quality contigs, the secondary/alternative contigs and high coverage contigs (e.g., putative organelle genomes, plastid, mitochondrion). Contigs showing a very high coverage and forming loop were blasted against the GenBank Non-Redundant Protein Sequence database (NCBI-BLAST) to verify the match with organelle genome. One contig (ctg011470, ~140 Kbp) showed high homology with *Medicago sativa* chloroplast sequences and five contig (ctg008800, ctg009160, ctg009300, ctg009310, ctg011650) showed homology with mitochondrial sequences of different plant species (*Q. robur, Q. acutissima* and *Lithocarpus litseifolius*).

**Supplementary Figure 3.**
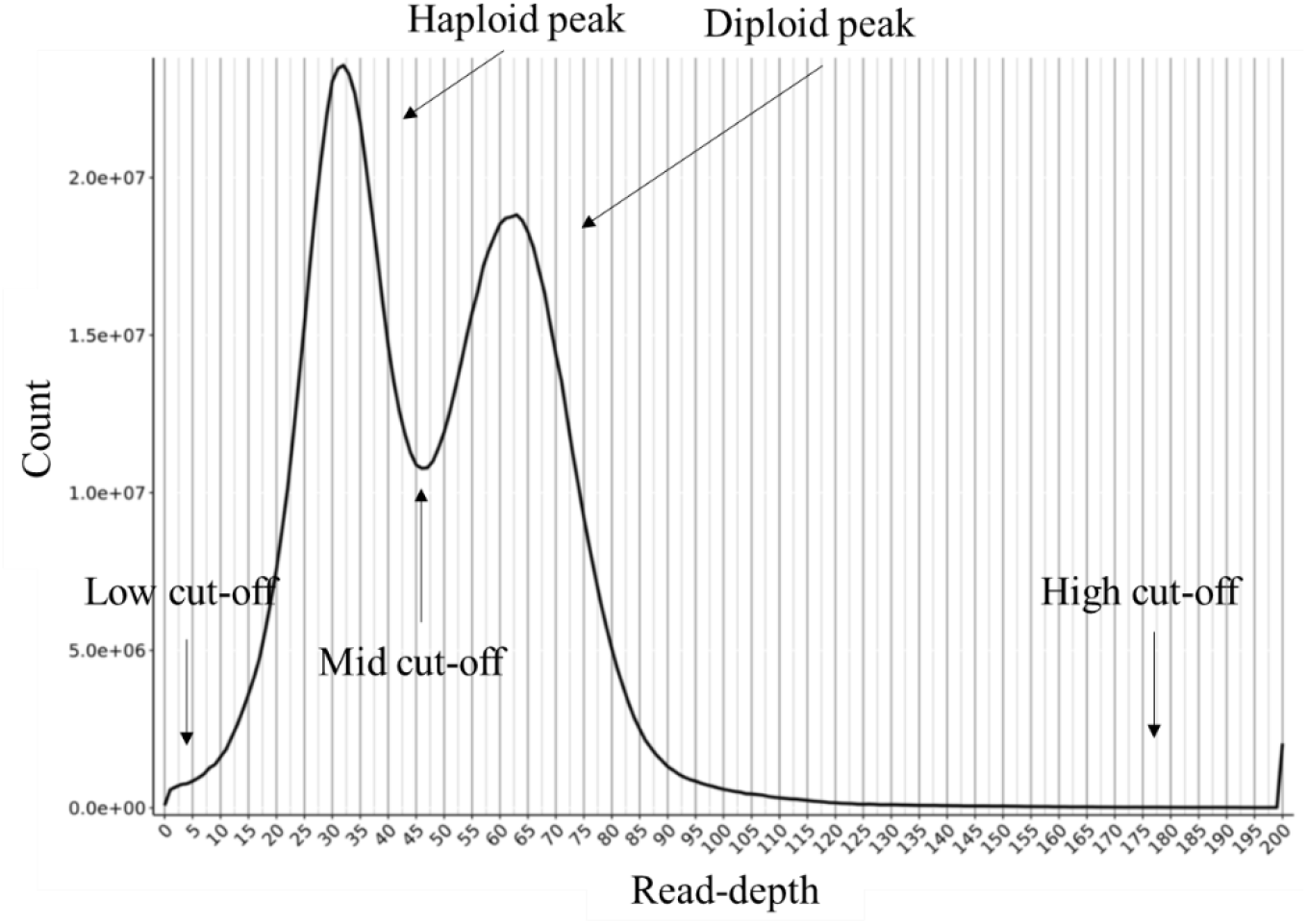
Distribution of read coverage. Arrows indicate the coverage points used to separate low, secondary, primary, and high repetitive contigs.

As a result, 530 contigs (total size 842.2 Mbp) were retained as primary contigs (Table 1), 616 (total size 157,3 Mbp) were flagged as secondary contigs and 20 (total size 1,1 Mbp) were flagged as artifacts (too low or too high coverage). The final assembly of the nuclear haploid genome had a total length of 842.2 Mbp, which is similar to a previous estimate using flow cytometry (935,0 Mbp) (Rey et al., 2019). The contig N50 had a size of 3.3 Mbp indicating a *de novo* assembly with high level of continuity. It is similar in size to other *Quercus* spp (Table 2).

The completeness of the final genome assembly (primary contigs) was evaluated using Benchmarking Universal Single-Copy Orthologs (BUSCO) version 5.0 (key parameters: default, -l viridiplantae_odb10) (Simão et al., 2015). Predicted gene models exhibited a completeness of 99% (Table 3), with a low percentage of duplicated (4%), fragmented (0.20%) and missing (0.80%) genes, indicating that the assembled genome has a high level of completeness.

**Table 3.** BUSCO assessment of the *Q. ilex* genome assembly (Database used: Viridiplantae).

| Gene number | Category | Percentage (%) |
| --- | --- | --- |
| 421 | Complete BUSCOs (C) | 99.00 |
| 403 | Complete and single-copy BUSCOs (S) | 94.80 |
| 18 | Complete and duplicated BUSCOs (D) | 4.20 |
| 1 | Fragmented BUSCOs (F) | 0.20 |
| 3 | Missing BUSCOs (M) | 0.80 |
| 425 | Total BUSCO groups searched |  |

Using a chromosome-based genome assembly of a close and syntenic species (*Q. lobata*, valley oak) NCBI ID=GCF_001633185.2, ValleyOak release 3.2), contigs were scaffolded at the chromosome level using RagTag (key parameters: scaffolding) (Alonge et al., 2019). The software performs a homology-based assembly. Briefly, each *Q. ilex* contig was mapped against the chromosomes of *Q. lobata* genome and scored for confidence in grouping, location, and orientation. As a result, 493 contigs, accounting for 835 Mbp (99% of the total assembly), were found to have some correspondence on *Q. lobata* chromosomes (Figure 2). Less than 1% of the sequences (37 contigs, size 6,69 Mbp) failed to be assigned. They were merged and placed in chromosome 0 (chr 0). Arbitrary gaps of 100 bp were added between consecutive contigs (517 gap sequences, 51.7 Kbp).

**Figure 2:**
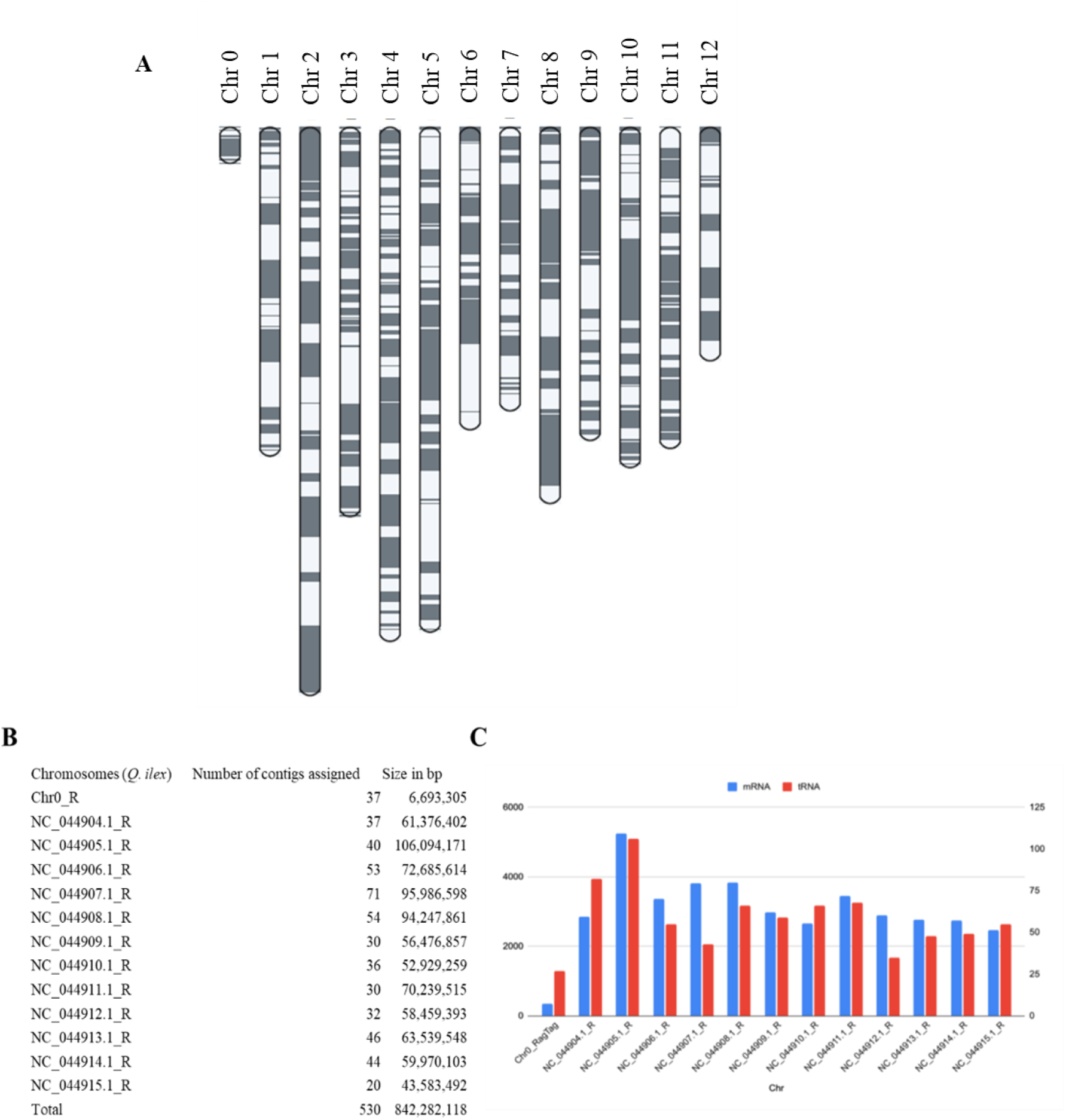
Homology-based *Q. ilex* chromosomes. (**A**) Alternance of grey and white bands represents consecutive contigs mapped to chromosomes. Chr stands for chromosome. (**B**) Number of *Q. ilex* contigs assigned to each chromosome and their relative size in bp. (**C**) mRNA (blue) and tRNA (red) distribution across *Q. ilex* chromosomes.

**Figure 4:**
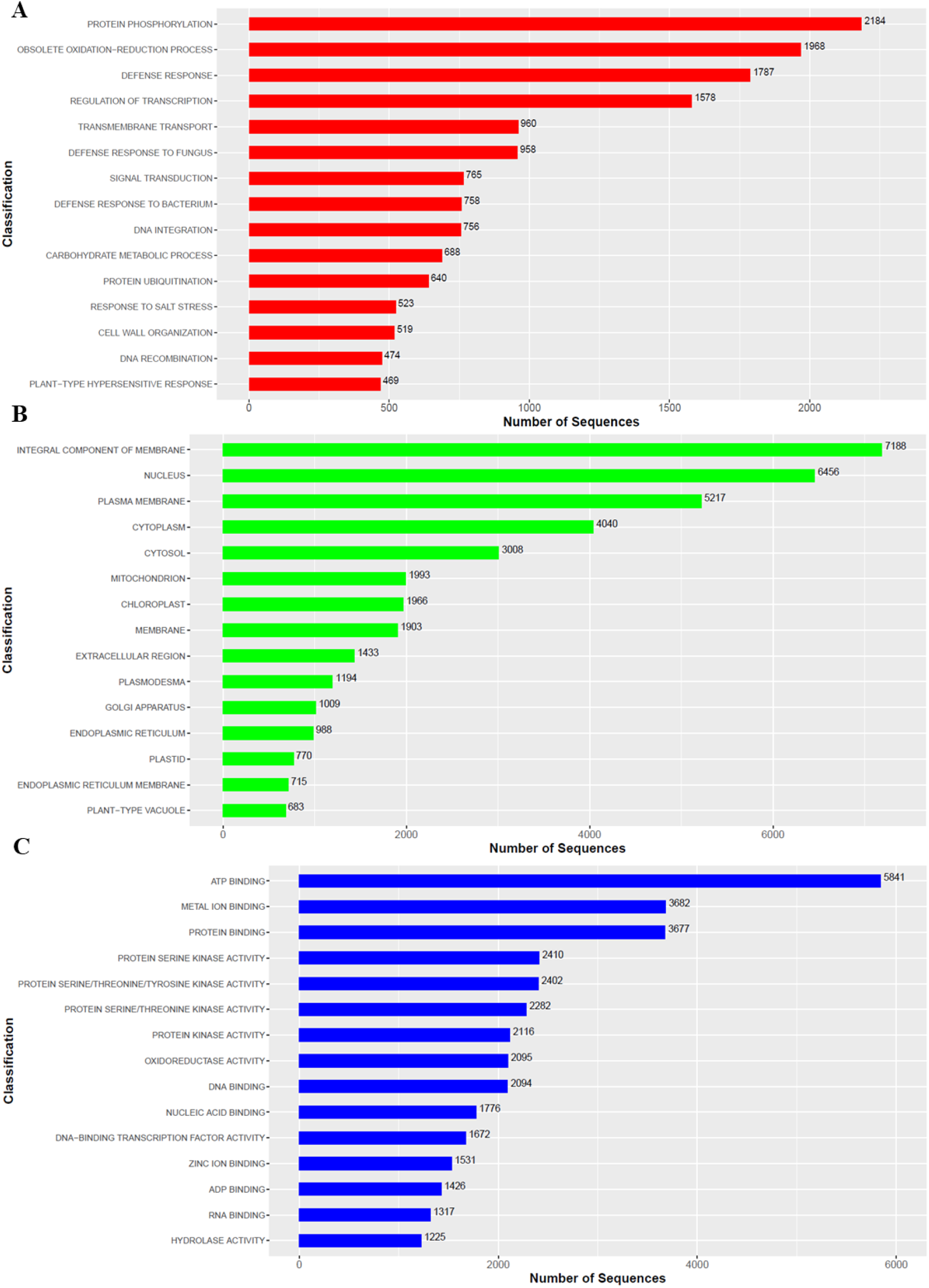
Number of *Quercus ilex* genes per GO categories: (**A**) Biological Processes, (**B**) Cellular Components, and (**C**) Molecular Functions.

**Figure 5:**
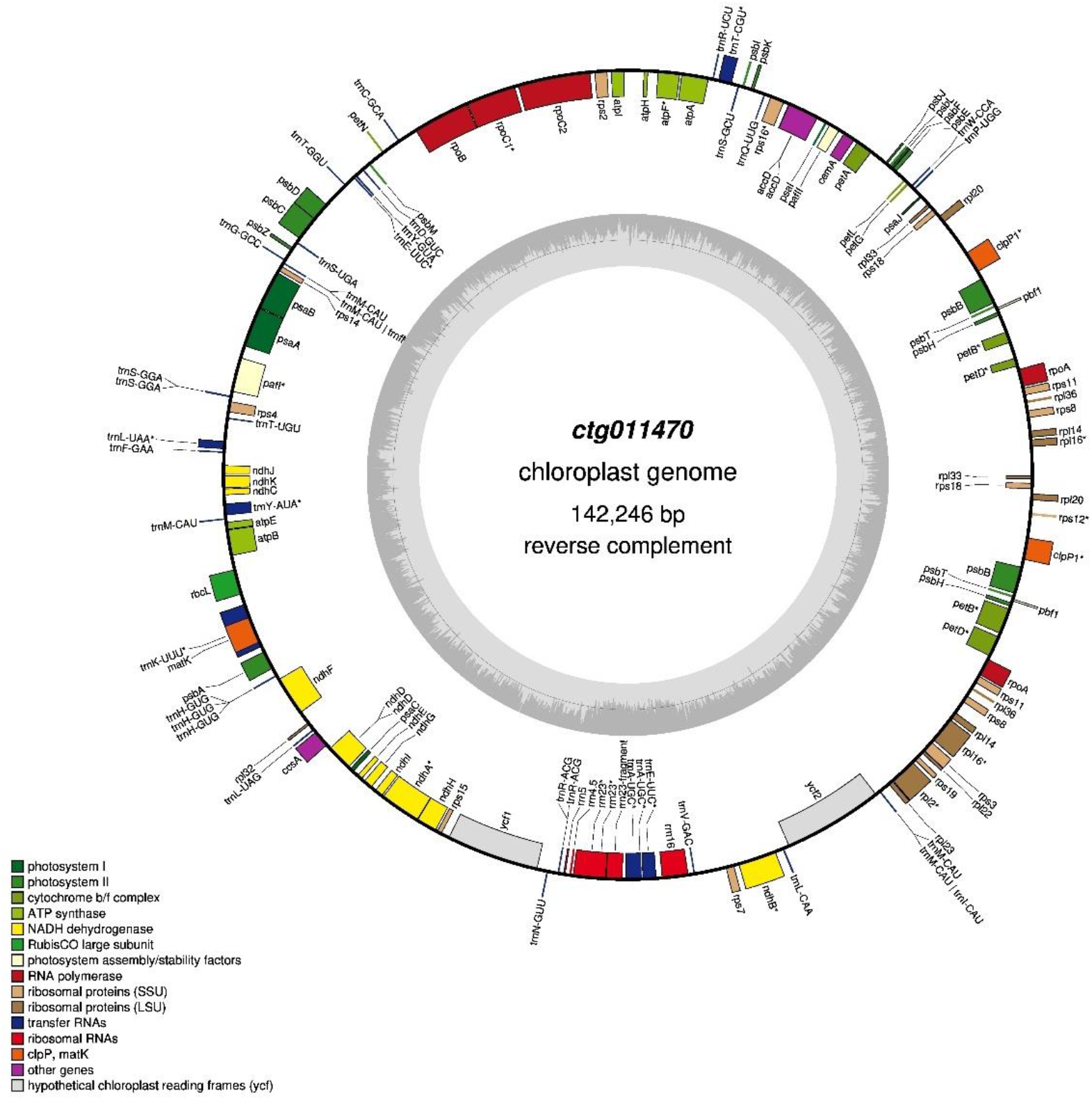
Gene map of the *Q. ilex* chloroplast genome. Genes shown outside and inside the outer circle are transcribed clockwise and counterclockwise, respectively. Genes belonging to different functional groups are represented in different colors. The darker and lighter gray areas in the inner circle indicate the GC and AT content, respectively.

### Repeat annotation

We generated a high-quality non-redundant transposable element (TE) library for *Q. ilex* genome using the Extensive De-novo TE Annotator (EDTA) pipeline (https://github.com/oushujun/EDTA) (Ou et al., 2019). The pipeline combines a suite of best-performing packages (LTR_FINDER, LTR_harvest, LTR_retriever, Generic Repeat Finder, TIR Learner, HelitronScanner and RepeatMasker), and it was designed to filter out false discoveries in raw TE candidates and generate high-quality non-redundant TE libraries. The inbuilt package RepeatModeler version 2.0.1 (key parameters: default) (Bedell et al., 2000) identifies any remaining TEs which might have been overlooked by the EDTA algorithm (--sensitive 1) and was also used in the workflow. TE identification was performed using RepeatMasker version 1.332 (key parameters: -s -nolow -norna -nois -e rmblast -gff) (Bedell et al., 2000) utilizing the NCBI/RMBLAST version 2.6.0+ search engine. Repeat sequences in the *Q. ilex* genome assembly accounted for 53.3% (448,68 Mb) of the total genome length (Table 4). LTR Gypsy and Copia were the most abundant repeat types (14.37% and 11.58%, respectively) (Han et al., 2022). The proportion of repeat elements in *Q. ilex* genome was similar to other *Quercus* spp. (Table 2).

**Table 4.**
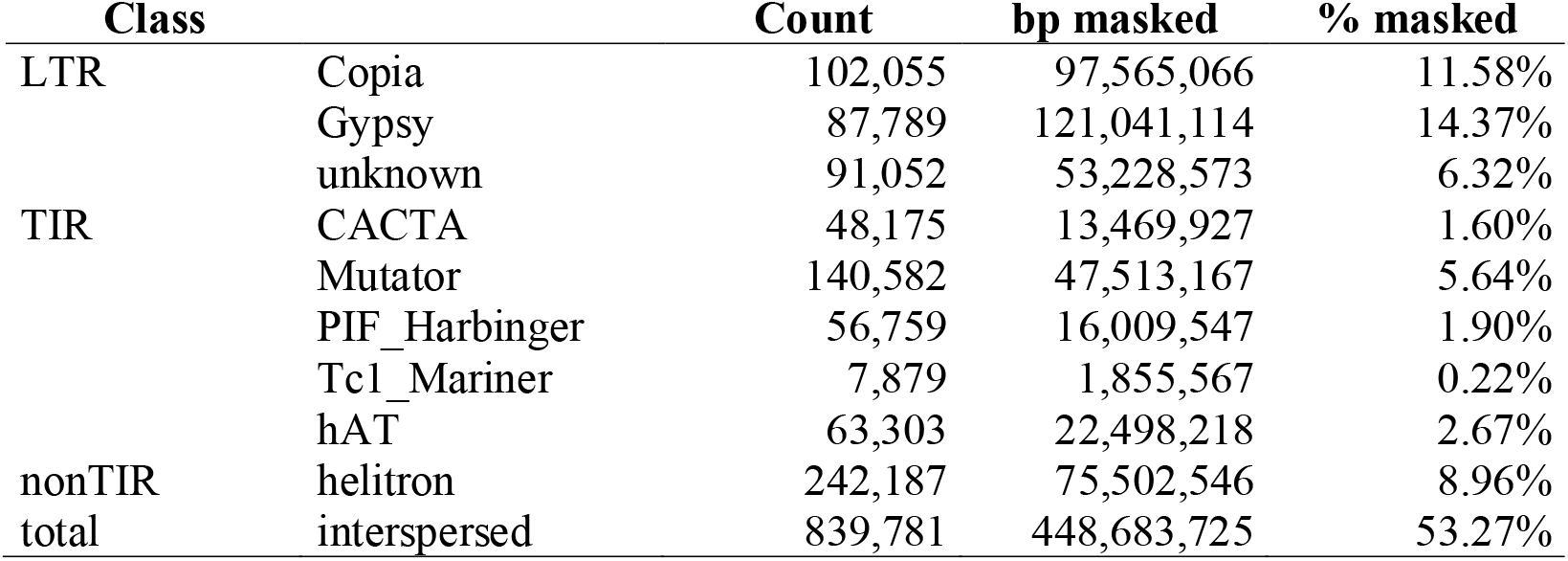
Whole-genome TE annotation summary statistics obtained through the EDTA pipeline

### Gene prediction and annotation

The annotation pipeline integrated different tasks starting from the fetching of raw data from public and/or private repositories (ESTs, protein collections, RNA-Seq datasets and other relevant sequences) to sequence alignment, prediction of gene models, and their functional annotations (Supplementary Figure 5).

**Supplementary Figure 5.**
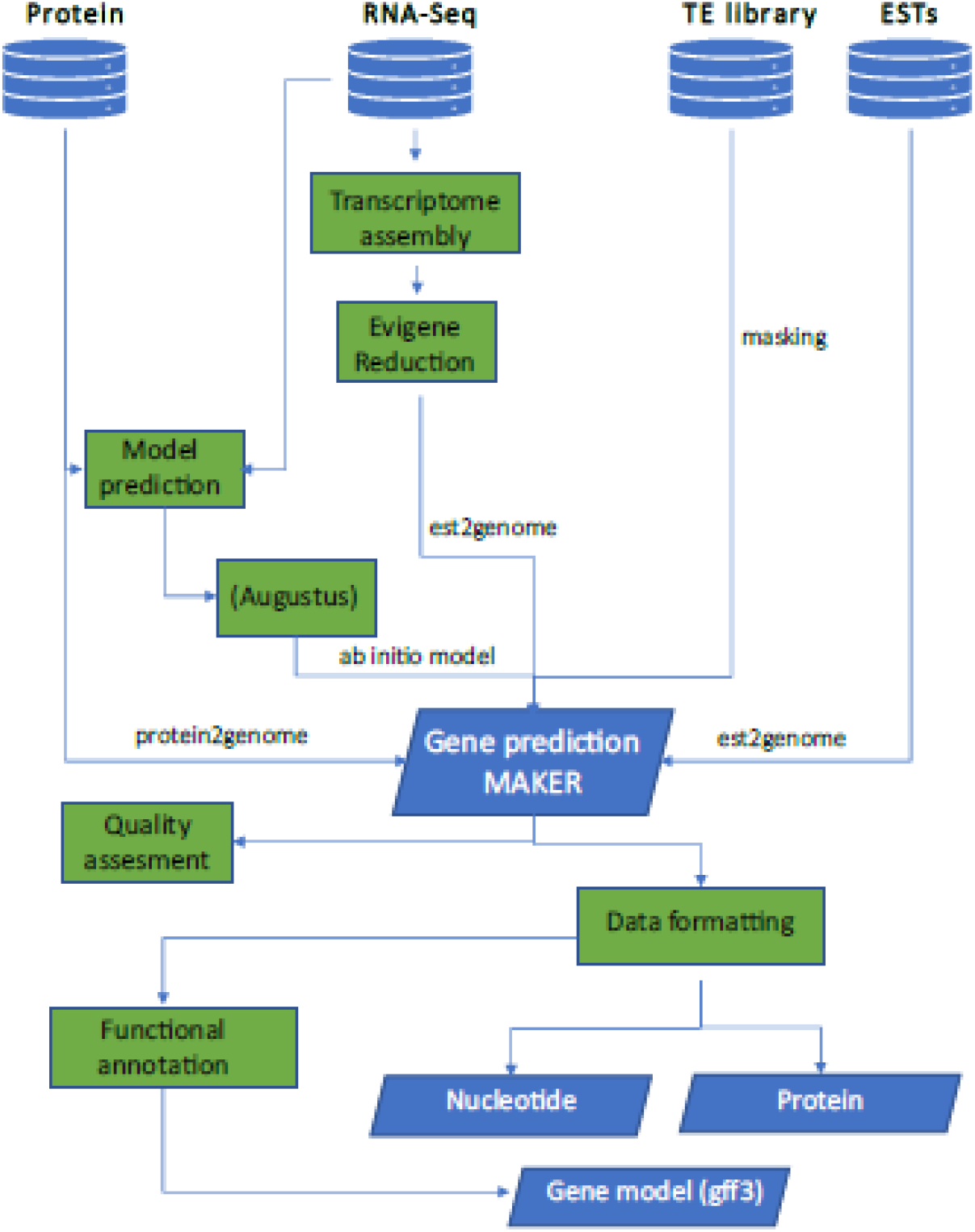
The genome annotation workflow

We used MAKER2 (key parameters: default) (Campbell et al., 2014) as the core module for structural genome annotation. The software synthesizes a final annotation based on quality evidence-values. Here, evidence was based on transcripts previously produced using RNA-Seq datasets (from NCBI dataset: SRR11050903 and SRX2993508) generated for *Q. ilex* by our research group (Guerrero-Sánchez et al., 2017, 2019, 2021), RNA-Seq dataset generated by Madritsch et al., (2018) from *Q. ilex* seedling leaves under drought conditions (from NCBI dataset: SRX4725055 and SRX4725058), 22,000 reviewed proteins from UniprotKb (taxa rosids), two sets of proteins of closed related species (*Q. lobata* and *Juglans regia*) (from NCBI dataset: GCF_001633185.2 in *Q. lobata* and GCA_001411555.2 in *J. regia*), EST collection of *Q. ilex* (download from NCBI) and the curated TE library produced by the EDTA pipeline generated in this work. The final gene set accounted for 39,443 protein-coding genes (Table 5; Supplementary Table 2). Also, a total of 759 tRNA were predicted (Supplementary Table 3) by using tRNAscan-SE (key parameter: default) (Lowe et al., 1997). The distribution of mRNA and transfer RNAs (tRNA) across chromosomes is reported in Figure 2C. Other *Quercus* spp. also showed similar number of protein-coding genes (Table 2).

**Supplementary Table 2.** List of protein-coding genes annotated in *Q. ilex* genome.

**Table 5.** Gene annotation summary for the *Q. ilex* genome.

|  |  |
| --- | --- |
| Number of gene/mRNA | 39,443 |
| Number of mRNAs with UTR both sides | 18,255 |
| Number of mRNAs with at least one UTR | 25,862 |
| Number of CDS | 39,443 |
| Number of exón | 192,536 |
| Number of five_prime_UTR | 20,321 |
| Number of three_prime_UTR | 23,796 |
| mean exons per mRNA | 4.9 |
| mean five_prime_UTRs per mRNA | 0.5 |
| mean three_prime_utrs per mRNA | 0.6 |
| Total gene/mRNA length (bp) | 156,941,371 |
| Total CDS length (bp) | 46,744,435 |
| Total exon length (bp) | 59,868,779 |
| Total five_prime_UTR length (bp) | 4,337,145 |
| Total three_prime_UTR length (bp) | 8,787,199 |
| mean gene/mRNA length (bp) | 3,978 |
| mean CDS length (bp) | 1,185 |
| mean exon length (bp) | 310 |
| mean five_prime_UTR length (bp) | 213 |
| mean three_prime_UTR length (bp) | 369 |
| % of genome covered by gene | 18.5 |
| % of genome covered by mRNA | 18.5 |
| % of genome covered by CDS | 5.5 |
| % of genome covered by exon | 7 |
| % of genome covered by five_prime_UTR | 0.5 |
| % of genome covered by three_prime_UTR | 1 |

**Supplementary Table 3.**
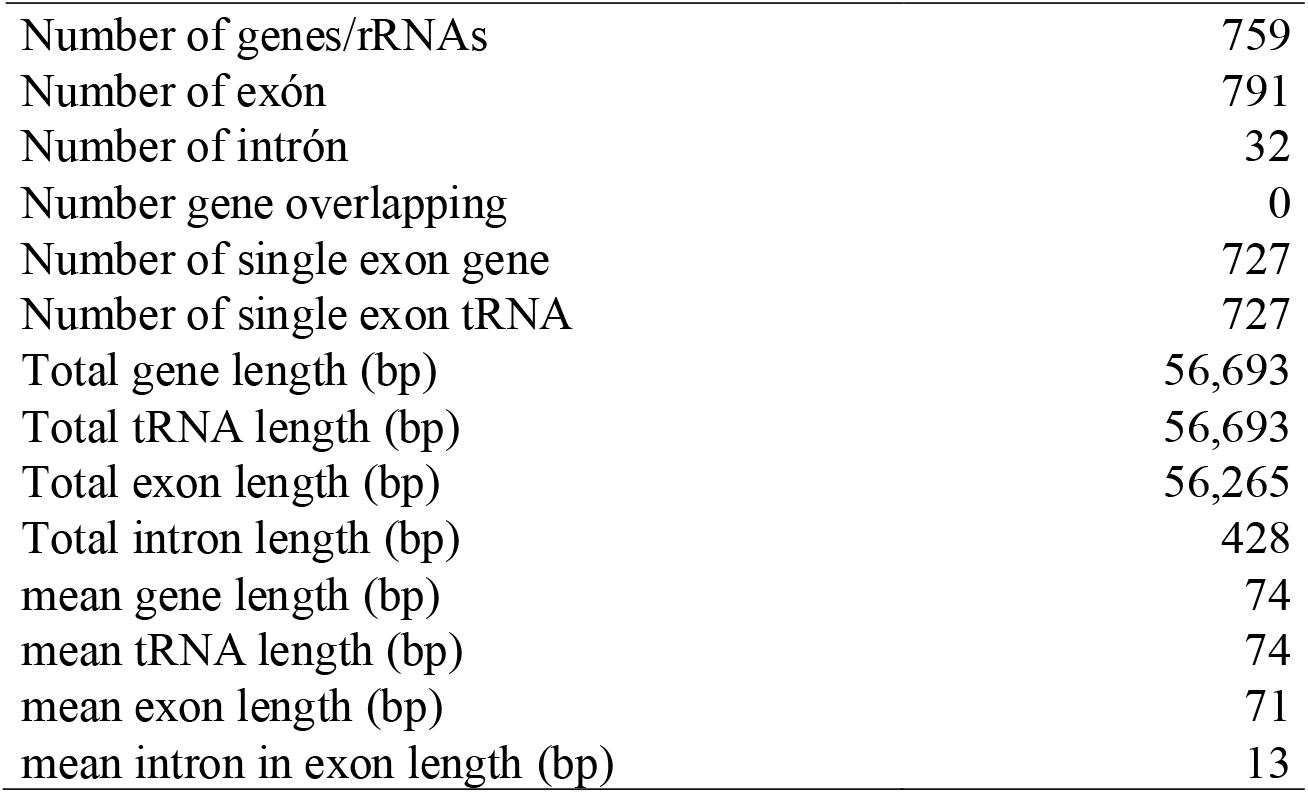
Annotation of tRNA for the *Q. ilex* assembled genome.

|  |  |
| --- | --- |
| Number of genes/rRNAs | 759 |
| Number of exón | 791 |
| Number of intrón | 32 |
| Number gene overlapping | 0 |
| Number of single exon gene | 727 |
| Number of single exon tRNA | 727 |
| Total gene length (bp) | 56,693 |
| Total tRNA length (bp) | 56,693 |
| Total exon length (bp) | 56,265 |
| Total intron length (bp) | 428 |
| mean gene length (bp) | 74 |
| mean tRNA length (bp) | 74 |
| mean exon length (bp) | 71 |
| mean intron in exon length (bp) | 13 |

A final assessment of the mRNA sequence was evaluated using BUSCO (Supplementary Table 4). Results showed a completeness > 98% with a high degree of single copy genes (>95%) and a low percentage of duplication (3.1%).

**Supplementary Table 4.** BUSCO assessment of the *Q. ilex* genome annotation.

| Gene number | Category | Percentage |
| --- | --- | --- |
| 418 | Complete BUSCOs (C) | 98.40% |
| 405 | Complete and single-copy BUSCOs (S) | 95.30% |
| 13 | Complete and duplicated BUSCOs (D) | 3.10% |
| 5 | Fragmented BUSCOs (F) | 1.20% |
| 2 | Missing BUSCOs (M) | 0.40% |
| 425 | Total BUSCO groups searched |  |

### Functional annotation

To assign functional descriptions, Gene Ontology (GO) terms and Kyoto Encyclopedia of Genes and Genomes (KEGG) pathway information to the gene models, sequences (transcripts/proteins) were functionally annotated with TRINOTATE version 2.0 (key parameters: protDB=uniprot_3745 (Viridiplantae), transdecoder + diamond + pfam) (https://github.com/Trinotate/Trinotate.github.io/blob/master/index.asciidoc). Of the total number of 39,443 genes, 35,258 (89.4%), 29,507 (74,8%), and 9,555 (24.2%) were identified with functional descriptions, GO terms, and path maps (KEGG), respectively (Supplementary Table 5). The remaining 4,185 (10.6%) genes were described without annotation. The GO representation of the whole gene set according to biological processes, cellular components, and molecular functions are summarized in Figure 3.

### Assembly of chloroplast genome

Among all thee contigs, one (ctg011470, 142.3 Kbp) matched chloroplast sequences in NCBI (best match “*Medicago sativa* chloroplast, complete genome”, identity 99%). Chloroplast annotation was performed using GeSeq tool (key paratemers: default) (Tillich et al., 2017). GeSeq analyses input sequence by comparing it against reference databases using BLAT (Kent, 2002). We used additional tools (ARAGORN, tRNA-scan, Chloë version 0.1.0) to annotate tRNA, rRNA and CDS, and OrganellarGenomeDRAW (Greiner et al., 2019) was used to convert annotations into a circular graphical map. The final chloroplast assembly was 142.3 Kbp, GC content of 33.5%, and 149 annotated genes of which 92 were protein-coding genes, 51 were rRNA and 6 were rRNA. Similar results have been previously described for *Q. robur* ‘Fastigiata’ and *Q. variabilis*, with chloroplast sizes of 161.2 Kbp (Feng et al., 2020) and 160.8 Kbp (Zeng et al., 2019), respectively.

## Conclusions

We provide a *de novo* genome assembly of *Q. ilex* based on long reads generated by SMRT technology from the PacBio platform. The first draft of the assembled genome was approximately 842.2 Mb in size, with high heterozygosity (1.93%), a high level of continuity with a contig N50 of 3.3 Mbp and completeness identifying 418 out of 425 annotated genes. Out of 39,443 genes, 35,258 had functional descriptions, 29,507 included GO terms and 9,555 were identified in a path map. Repeat sequences accounted for 53.27% of the genome, with LTR Gypsy and Copia being the most abundant (25,95%). A complete (i.e. circular without gaps) chloroplast genome was assembled with a size of 142.3 Kbp. The available first draft of the assembled genome, together with previously published transcriptomics, proteomics, and metabolomics datasets (San-Eufrasio et al., 2021; Guerrero-Sánchez et al., 2021; López-Hidalgo et al., 2021; Tienda-Parrilla et al., 2021; Maldonado-Alconada et al., 2022), provide valuable information to future studies on comparative genomics, and studies of breeding for disease resistance and nutritional quality in acorns of *Q. ilex*.

## Availability of Supporting Data and Material

The raw genomic sequences were deposited in the NCBI SRA database under the BioProject accession number PRJNA687489.

## Acknowledgement

We thank Bruce Kingham and Olga Shevchenko at the Sequencing and Genotyping Center at the University of Delaware DNA for assistance and sequencing of the raw data of the *Q. ilex* genome.

## Authors’ s Contributions

J.V.J.N. performed sample collection. M.D.R. extracted genomic DNA. V.R. conducted the whole genome sequencing analysis. M.L.O performed the microsatellite analysis. M.D.R., V.M.G.S., R.C., M.A.C., A.R.F., R. B., and J.V.J.N. were involved in data interpretation. M.D.R. drafted the original manuscript. M.D.R., R.C., and J.V.J.N. finalized the manuscript. All authors read and approved the final manuscript.

## References

Alonge M, Soyk S, Ramakrishnan S, et al. RaGOO: fast and accurate reference-guided scaffolding of draft genomes. Genome Biol 2019;20:1–17.

Backs JR, Ashley MV. Quercus Genetics: Insights into the Past, Present, and Future of Oaks. Forests 2021;12:1628.

Barbeta A, Peñuelas J. Sequence of plant responses to droughts of different timescales: lessons from holm oak (Quercus ilex) forests. Plant Ecol Divers 2016;9:321–338.

Bedell JA, Korf I, Gish W. MaskerAid: a performance enhance-ment to RepeatMasker. Bioinformatics 2000;16:1040–1041.

Brasier CM. Oak tree mortality in Iberia. Nature 1992;360:539–539.

Burgarella C, Lorenzo Z, Jabbour-Zahab R, et al. Detection of hybrids in nature: application to oaks (Quercus suber and Q. ilex). Heredity 2009;102:442–452.

Camilo-Alves CSP, Clara MIE, Ribeiro NMCA. Decline of Mediterranean oak trees and its association with Phytophthora cinnamomi: a review. Eur J For Res 2013;132:411–432.

Campbell MS, Law M, Holt C, et al. MAKER-P: a tool kit for the rapid creation, management, and quality control of plant genome annotations. Plant Physiol 2014;164:513–524.

Cortés AJ, Restrepo-Montoya M, Bedoya-Canas LE. Modern strategies to assess and breed forest tree adaptation to changing climate. Front Plant Sci 2020;11:583323.

De Rigo D, Caudullo G. Quercus ilex in Europe: distribution, habitat, usage and threats. European atlas of forest tree species 2016;152–153.

Díaz-Caro C, García-Torres S, Elghannam A, et al. Is production system a relevant attribute in consumers’ food preferences? The case of Iberian dry-cured ham in Spain. Meat Sci 2019; 158:107908.

Escandón M, Castillejo-Sánchez MA, Jorrín-Novo JV, et al. Molecular research on stress responses in Quercus spp.: From classical biochemistry to systems biology through omics analysis. Forests 2021;12:364.

Feng L, Yang X, Jiao Q, et al., The complete chloroplast genome of Quercus robur ‘Fastigiata’. Mitochondrial DNA B: Resour 2020;5:129–130.

Fernández i Marti A, Romero-Rodríguez C, Navarro-Cerrillo RM, et al. Population genetic diversity of Quercus ilex subsp. ballota (Desf.) samp. reveals divergence in recent and evolutionary migration rates in the spanish dehesas. Forests 2018;9:337.

Greiner S, Lehwark P, Bock R. OrganellarGenomeDRAW (OGDRAW) version 1.3.1: expanded toolkit for the graphical visualization of organellar genomes. Nucleic Acids Res 2019;47:W59–W64.

Guerrero-Sanchez VM, Maldonado-Alconada AM, Amil-Ruiz F, et al. Holm oak (Quercus ilex) transcriptome. De novo sequencing and assembly analysis. Front Mol Biosci 2017;4:70.

Guerrero-Sanchez VM, Maldonado-Alconada AM, Amil-Ruiz F, Ion Torrent and lllumina, two complementary RNA-seq platforms for constructing the holm oak (Quercus ilex) transcriptome. PLoS One 2019;14:e0210356.

Guerrero-Sanchez VM, Castillejo MÁ, López-Hidalgo C, et al. Changes in the transcript and protein profiles of Quercus ilex seedlings in response to drought stress. J Proteom 2021;243:104263.

Han B, Wang L, Xian Y, et al. A chromosome-level genome assembly of the Chinese cork oak (Quercus variabilis). Front Plant Sci 2022;13.

Jombart T. adegenet: a R package for the multivariate analysis of genetic markers. Bioinformatics 2008;24:1403–1405.

Kajitani R, Toshimoto K, Noguchi H, et al. Efficient de novo assembly of highly heterozygous genomes from whole-genome shotgun short reads. Genome Res 2014;24:1384–1395.

Kent WJ. BLAT–the BLAST-like alignment tool. Genome Res 2002;12:656–664.

Laureano RG, García-Nogales A, Seco JI, et al. Plant maintenance and environmental stress. Summarising the effects of contrasting elevation, soil, and latitude on Quercus ilex respiration rates. Plant Soil 2016;409:389–403.

Leroy T, Plomión C, Kremer C. Oak symbolism in the light of genomics. New Phytol 2020;226:1012–1017.

López De Heredia U, Sanchez H, Soto A. Molecular evidence of bidirectional introgression between Quercus suber and Quercus ilex. IFOREST 2018;11:338.

López De Heredia U, Mora-Márquez F, Goicoechea PG, et al. ddRAD sequencing-based identification of genomic boundaries and permeability in Quercus ilex and Q. suber hybrids. Front Plant Sci 2020;1330.

López-Hidalgo C, Menéndez M, Jorrín-Novo JV. Phytochemical composition and variability in Quercus ilex acorn morphotypes as determined by NIRS and MS-based approaches. Food Chem 2021;338:127803.

Lowe TM, Eddy SR. tRNAscan-SE: a program for improved detection of transfer RNA genes in genomic sequence. Nucleic Acids Res 1997;25:955–964.

Madritsch S, Wischnitzki E, Kotradre P, et al. Elucidating drought stress tolerance in European oaks through cross-species transcriptomics. G3 (Bethesda, Md.) 2019;9:3181–3199.

Maldonado-Alconada AM, Castillejo MÁ, Rey MD, et al. Multiomics Molecular Research into the Recalcitrant and Orphan Quercus ilex Tree Species: Why, What for and How. Int J Mol Sci 2022;23:9980.

Murray MG, Thompson WF. Rapid isolation of high molecular weight plant DNA. Nucleic Acids Res 1980;8:4321–4326.

Muthuramalingam P, Jeyasri R, Rakkammal K, et al. Multi-Omics and Integrative Approach towards Understanding Salinity Tolerance in Rice: A Review. Biology 2022;11:1022.

Ou S, Su W, Liao Y, et al. Benchmarking transposable element annotation methods for creation of a streamlined, comprehensive pipeline. Genome Biol 2019;20:1–18.

Paajanen P, Kettleborough G, López-Girona E, et al. A critical comparison of technologies for a plant genome sequencing project. Gigascience 2019;8:giy163.

Plieninger T, Flinzberger L, Hetman M, et al. Dehesas as high nature value farming systems: a social-ecological synthesis of drivers, pressures, state, impacts, and responses. Ecol Soc 2021;26.

Plomión C, Aury J, Amselem J, et al. Decoding the oak genome: Public release of sequence data, assembly, annotation and publication strategies. Mol Ecol Resour 2016;16:254–265.

Plomión C, Aury, J, Amselem J, et al. Oak genome reveals facets of long lifespan. Nat Plants 2018;4:440–452.

Ramos AM, Usie A, Barbosa P, et al. The draft genome sequence of cork oak. Sci Data 2018;5:180069.

Rey MD, Castillejo MÁ, Sánchez-Lucas R, et al. Proteomics, holm oak (Quercus ilex L.) and other recalcitrant and orphan forest tree species: how do they see each other?. Int J Mol Sci 2019;20:692.

Roach MJ, Schmidt SA, Borneman AR Purge Haplotigs: allelic contig reassignment for third-gen diploid genome assemblies. BMC Bioinform 2018;19:1–10.

Roberts RJ, Carneiro MO, Schatz MC. The advantages of SMRT sequencing. Genome Biol 2013;14:405

Ruiz-Gómez F, Pérez-De-Luque A, Sánchez-Cuesta R, Differences in the response to acute drought and Phytophthora cinnamomi Rands infection in Quercus ilex L. seedlings. Forests 2018;9:634.

San-Eufrasio B, Sánchez-Lucas R, López-Hidalgo C, et al. Responses and differences in tolerance to water shortage under climatic dryness conditions in seedlings from Quercus spp. and Andalusian Q. ilex populations. Forests 2020;11:707.

San-Eufrasio B, Castillejo MÁ, Labella-Ortega M, Effect and response of Quercus ilex subsp. ballota [Desf.] Samp. seedlings from three contrasting Andalusian populations to individual and combined Phytophthora cinnamomi and drought stresses. Front Plant Sci 2021;12.

Simão FA, Waterhouse RM, Ioannidis P, et al. BUSCO: assessing genome assembly and annotation completeness with single-copy orthologs. Bioinformatics 2015;31:3210–3212.

Sork VL, Fitz-Gibbon S, Puiu D, et al. First draft assembly and annotation of the genome of a California Endemic oak Quercus lobata Née (Fagaceae). G3 Genes Genomes Genet 2016;6:3485–3495.

Sork VL, Cokus SJ, Fitz-Gibbon ST, et al. High-quality genome and methylomes illustrate features underlying evolutionary success of oaks. Nat commun 2022:13:1–15.

Soto A, Lorenzo Z, Gill L. Differences in fine-scale genetic structure and dispersal in Quercus ilex L. and Q. suber L.: Consequences for regeneration of mediterranean open woods. Heredity 2007;99:601–607.

Stocks JJ, Metheringham CL, Plumb WJ, et al. Genomic basis of European ash tree resistance to ash dieback fungus. Nat. Ecol. Evol. 2019;3:1686–1696.

Tillich M, Lehwark P, Pellizzer T, et al. GeSeq – versatile and accurate annotation of organelle genomes. Nucleic Acids Res 2017;45:W6–W11.

Toumi L, Lumaret R. Allozyme variation in cork oak (Quercus suber L.): the role of phylogeography and genetic introgression by other Mediterranean oak species and human activities. Theor Appl Genet 1998;97:647–656.

Valero-Galván J, Valledor L, Navarro-Cerrillo RM, et al. Studies of variability in Holm oak (Quercus ilex subsp. ballota [Desf.] Samp.) through acorn protein profile analysis. J Proteom 2011;74:1244–1255.

Vinha AF, Costa ASG, Barreira JC, et al. Chemical and antioxidant profiles of acorn tissues from Quercus spp.: Potential as new industrial raw materials. Ind Crops Prod 2016;94:143–151.

Xia EH, Zhang HB, Sheng J, et al. The tea tree genome provides insights into tea flavor and independent evolution of caffeine biosynthesis. Mol Plant 2017;10:866–877.

Zhao D, Hamilton JP, Pham GM, et al. De novo genome assembly of Camptotheca acuminata, a natural source of the anti-cancer compound camptothecin. GigaScience 2017;6:gix065.

Zeng QM, Liu B, Lin RQ, et al. The complete chloroplast genome sequence of Quercus gilva (Fagaceae). Mitochondrial DNA B: Resour 2019;4:2493–2494.

